## Supplemental Information for "ASPSCR1-TFE3 reprograms transcription by organizing enhancer loops around hexameric VCP/p97"

### Supplementary Information

| Supplementary Information | pages |
|---|---|
| Extended Data Figures with Legends | 2 - 15 |
| Supplementary Tables | 16 - 23 |

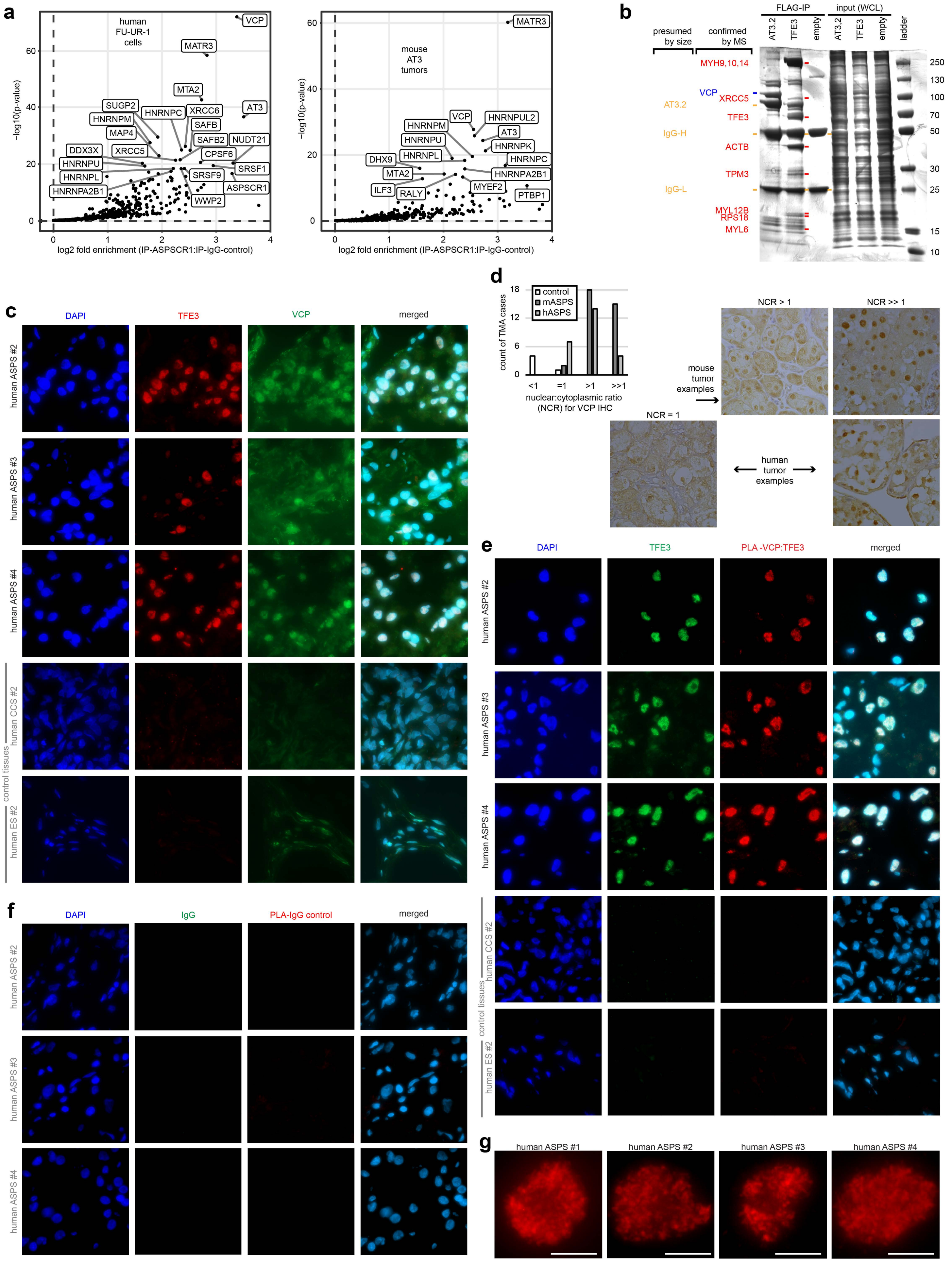
a
human
FU-UR-1
cells
VCP
MATR3
MTA2
SUGP2
HNRNPM
HNRNPC
XRCC6
AT3
SAFB
MAP4
SAFB2
NUDT21
DDX3X
XRCC5
CPSF6
HNRNPU
SRSF1
HNRNPL
SRSF9
ASPSCR1
HNRNPA2B1
WWP2
−log10(p-value)
log2 fold enrichment (IP-ASPSCR1:IP-IgG-control)
mouse
AT3
tumors
MATR3
VCP
HNRNPUL2
HNRNPM
AT3
HNRNPU
HNRNPK
HNRNPL
HNRNPC
DHX9
MTA2
HNRNPA2B1
MYEF2
ILF3
RALY
PTBP1
b
presumed
by size
confirmed
by MS
FLAG-IP
input (WCL)
AT3.2
TFE3
empty
ladder
MYH9,10,14
VCP
XRCC5
AT3.2
TFE3
IgG-H
ACTB
TPM3
IgG-L
MYL12B
RPS18
MYL6
250
130
100
70
50
30
25
15
10
c
DAPI
TFE3
VCP
merged
human ASPS #2
human ASPS #3
human ASPS #4
human CCS #2
human ES #2
control tissues
d
control
mASPS
hASPS
count of TMA cases
nuclear:cytoplasmic ratio
(NCR) for VCP IHC
NCR > 1
NCR >> 1
NCR = 1
mouse
tumor
examples
human
tumor
examples
e
DAPI
TFE3
PLA -VCP:TFE3
merged
f
DAPI
IgG
PLA-IgG control
merged
g
human ASPS #1
human ASPS #2
human ASPS #3
human ASPS #4

**Extended Data Figure 1. ASPSCR1-TFE3 interacts with VCP/p97 in the nucleus of ASPS and RCC tumor cells through its ASPSCR1 portion.** (**a**) Identification of the proteins enriched in both human FU-UR-1 cells and mouse AT3-induced tumor tissues by IP-MS of anti-ASPSCR1 IPs from nuclear lysates. (**b**) Coomassie stain of FLAG-IP and whole cell lysate (WCL) input from HEK transfected with TFE3, AT3.2, or empty vector, indicating the MS-identified bands that were visually discrepant between the two IP baits. (**c**) Fluorescence photomicrographs of 5 human tumors, three additional ASPS tumors and two controls with an additional clear cell sarcoma (CCS) and an additional Ewing sarcomas (ES), stained with DAPI, anti-TFE3 (to detect AT3) or anti-VCP immunofluorescent antibodies. (Each image panel represents 100µm square). (**d**) Bar chart of the number of cases staining for variable ratios of nuclear-to-cytoplasmic (NCRs) staining for VCP immunohistochemistry (IHC) on two tissue microarrays (TMAs) for ASPS tumors, one from human tumors and another from mouse tumors. Representative photomicrographs of different ratios of staining from the mouse and human TMAs. (Each image panel represents 100µm square). (**e**) Fluorescence photomicrographs demonstrating the proximity ligation assay (PLA) for the interaction between anti-TFE3 and anti-VCP antibodies in the same three ASPS tumors and two control human tumor tissues as in C. (Each image panel represents 100µm square). (**f**) Fluorescence photomicrographs demonstrating the negative control PLA for the interaction between control antibodies in the three additional human ASPS tumor tissues from **Figure 1**. (Each image panel represents 100µm square). (**g**) Higher magnification photomicrographs of single nuclei from each ASPS tumor, demonstrating the PLA signal character (magnification bars = 5µm).

Extended Data Figure 2

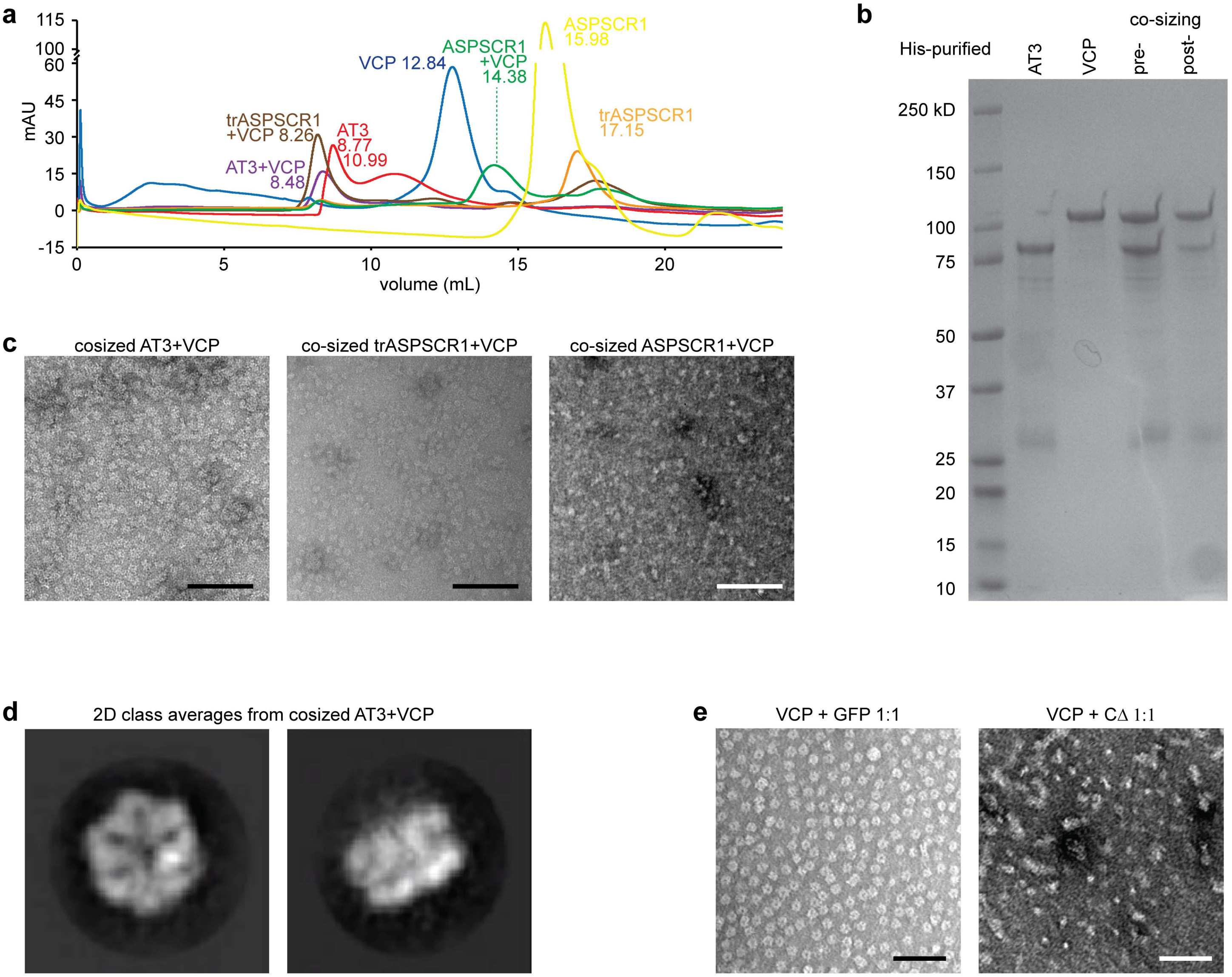


**Extended Data Figure 2. ASPSCR1-TFE3 interacts with hexameric assemblies of VCP/p97.** (**a**) Co-sizing size exclusion chromatography of each individual purified recombinant protein (ASPSCR1, trASPSCR1, AT3), as well as each after combining with VCP. (**b**) Coomassie stained SDS-PAGE of AT3 and VCP proteins before and after co-sizing and selection of the fraction representing the combination peak (8.49 mL in panel A). (**c**) Negative stain TEM of co-sized fractions of recombinant AT3, trASPSCR1, and ASPSCR1, each with VCP (scale bars = 100nm). (**d**) 2D class averages showing a top or pore view (1,406 particles) and side view (2,301 particles) of VCP hexamers, identified on co-sizing with AT3, but not ASPSCR1 full length (box length and width = 25 nm). (**e**) Negative stain TEM of VCP combined with equimolar GFP control or CΔ (scale bars = 100nm).

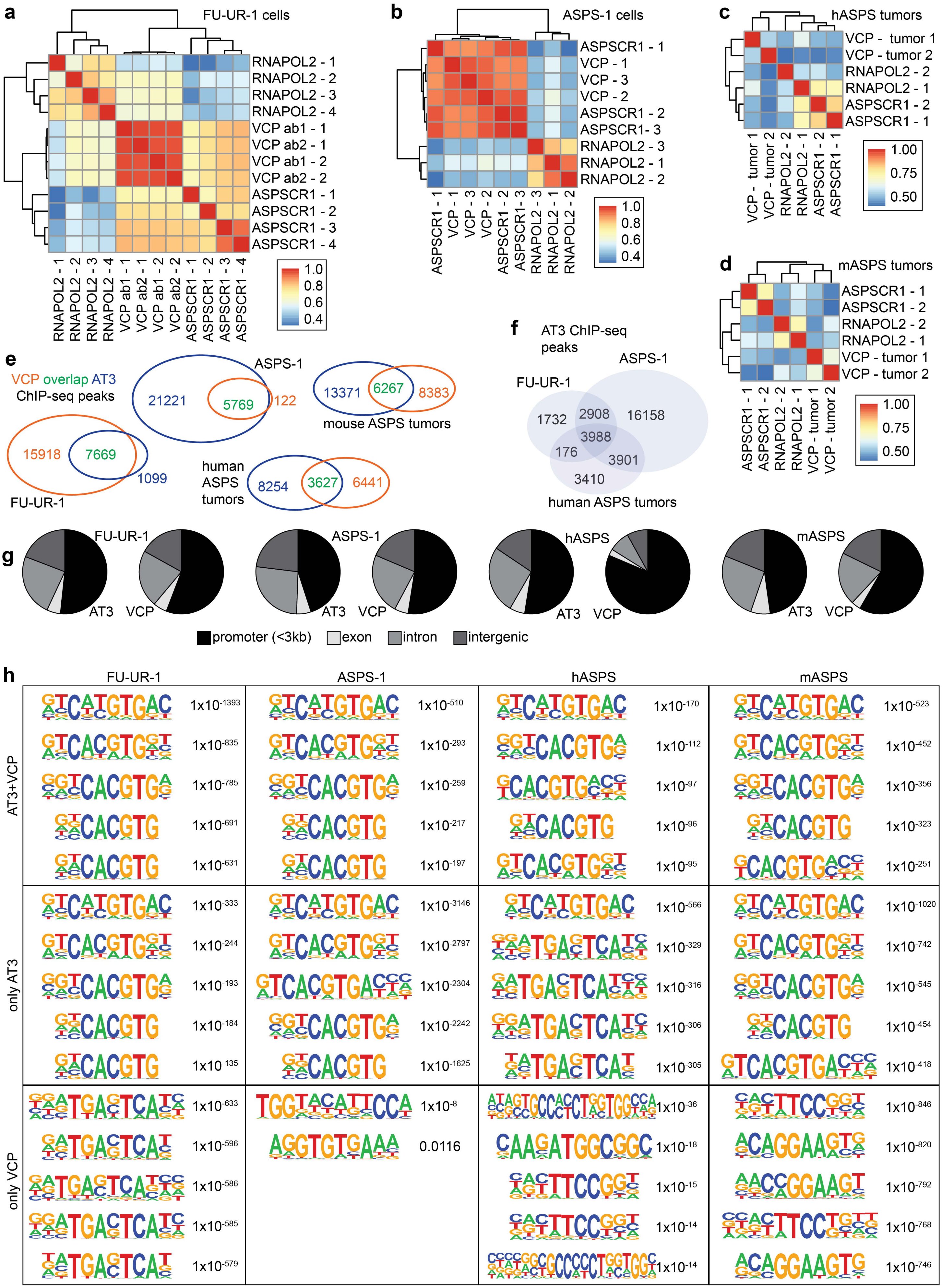
a
FU-UR-1 cells
RNAPOL2 - 1
RNAPOL2 - 2
RNAPOL2 - 3
RNAPOL2 - 4
VCP ab1 - 1
VCP ab2 - 1
VCP ab1 - 2
VCP ab2 - 2
ASPSCR1 - 1
ASPSCR1 - 2
ASPSCR1 - 3
ASPSCR1 - 4
1.0
0.8
0.6
0.4
b
ASPS-1 cells
ASPSCR1 - 1
VCP - 1
VCP - 3
VCP - 2
ASPSCR1 - 2
ASPSCR1- 3
RNAPOL2 - 3
RNAPOL2 - 1
RNAPOL2 - 2
1.0
0.8
0.6
0.4
c
hASPS tumors
VCP - tumor 1
VCP - tumor 2
RNAPOL2 - 2
RNAPOL2 - 1
ASPSCR1 - 2
ASPSCR1 - 1
1.00
0.75
0.50
d
mASPS tumors
ASPSCR1 - 1
ASPSCR1 - 2
RNAPOL2 - 2
RNAPOL2 - 1
VCP - tumor 1
VCP - tumor 2
1.00
0.75
0.50
e
VCP overlap AT3
ChIP-seq peaks
ASPS-1
21221
5769
122
13371
6267
8383
mouse ASPS tumors
15918
7669
1099
FU-UR-1
human ASPS tumors
8254
3627
6441
f
AT3 ChIP-seq peaks
ASPS-1
FU-UR-1
1732
2908
16158
3988
176
3901
3410
human ASPS tumors
g
FU-UR-1
ASPS-1
hASPS
mASPS
AT3
VCP
promoter (<3kb)
exon
intron
intergenic
h
FU-UR-1
ASPS-1
hASPS
mASPS
AT3+VCP
only AT3
only VCP
1x10-1393
1x10-835
1x10-785
1x10-691
1x10-631
1x10-510
1x10-293
1x10-259
1x10-217
1x10-197
1x10-170
1x10-112
1x10-97
1x10-96
1x10-95
1x10-523
1x10-452
1x10-356
1x10-323
1x10-251
1x10-333
1x10-244
1x10-193
1x10-184
1x10-135
1x10-3146
1x10-2797
1x10-2304
1x10-2242
1x10-1625
1x10-566
1x10-329
1x10-316
1x10-306
1x10-305
1x10-1020
1x10-742
1x10-545
1x10-454
1x10-418
1x10-633
1x10-596
1x10-586
1x10-585
1x10-579
1x10-8
0.0116
1x10-36
1x10-18
1x10-15
1x10-14
1x10-14
1x10-846
1x10-820
1x10-792
1x10-768
1x10-746

**Extended Data Figure 3. ASPSCR1-TFE3 and VCP interact on chromatin by ChIP-seq.** (**a**) Pearson correlation heatmap of biological replicates of ChIP-seq using antibodies against RNAPOL2, VCP, and ASPSCR1 amino terminus for AT3 in FU-UR-1 cells, (**b**) ASPS-1 cells, (**c**) 2 human ASPS tumors, and (**d**) two mouse AT3-initiated ASPS tumors. (**e**) Venn diagrams indicating the overlap of called peaks for AT3 with VCP in different models. (**f**) Venn diagrams indicating the overlap of called peaks for AT3 between the two human cell lines and human ASPS tumors. (**g**) Annotation pie charts for AT3 and VCP called peaks genome-wide with respect to genes between the 4 different systems. (**h**) Representation of the 5 most common known motifs in each subset of peaks from each group in Fig. 3a, peaks called for both AT3 and VCP, those called only for AT3, and those called only for VCP, with listed p-values.

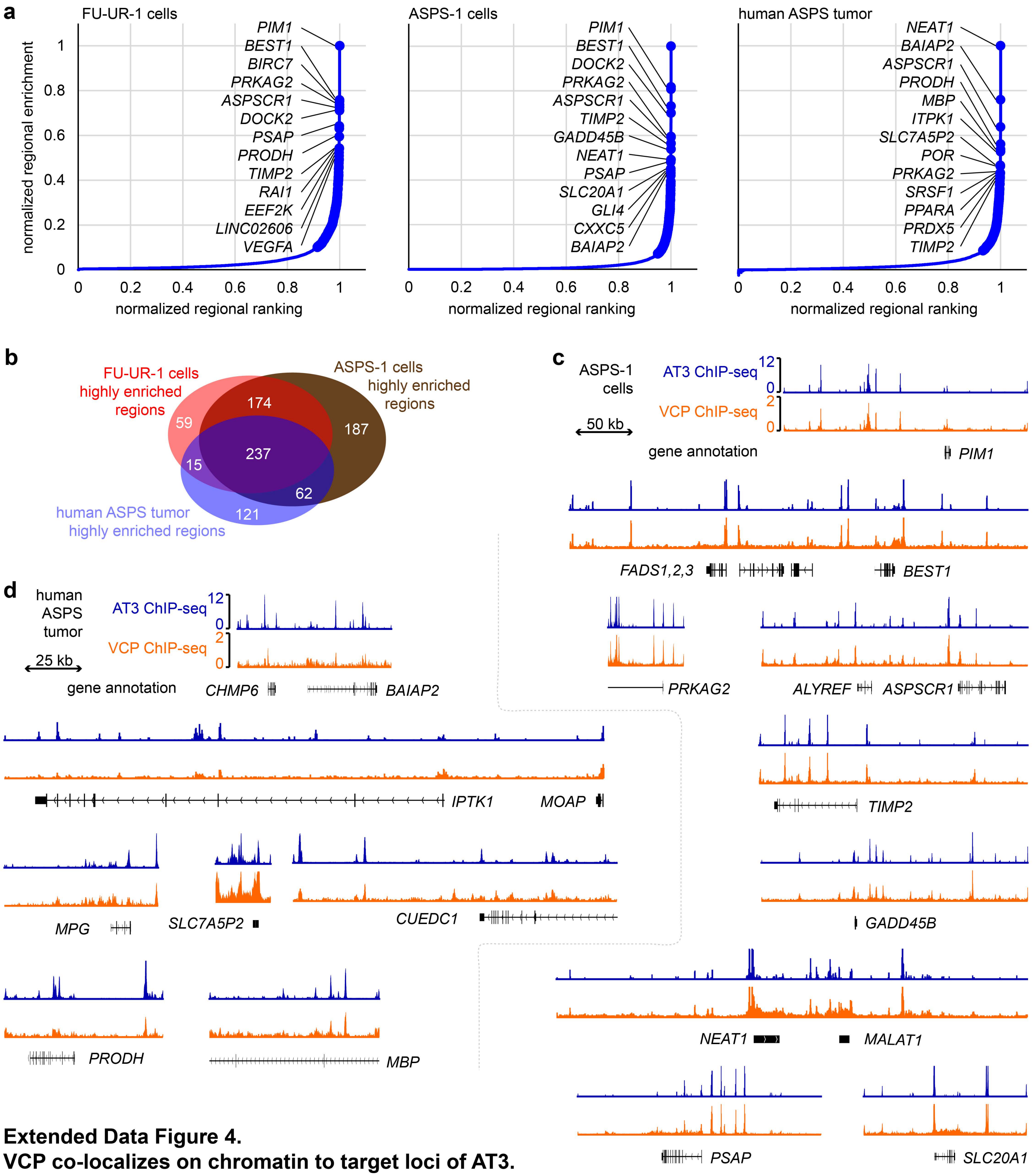


**Extended Data Figure 4.**
**VCP co-localizes on chromatin to target loci of AT3.**
(**a**) Plots of the normalized enrichment for AT3 ChIP-seq over the normalized ranking of enrichment for AT3 ChIP-seq in regions stitched together by stringently called peaks within a 40kb gap of each other for FU-UR-1 cells, ASPS-1 cells, and human ASPS tumor samples. All regions above the slope inflection point of 1 were designated highly enriched regions (and represented approximately the top 5 to 6% of regions). (**b**) Venn diagram of the location-overlapping highly enriched AT3 regions in human FU-UR-1 cells, ASPS-1 cells, and human ASPS tumors. (**c**) Representative tracks from highly enriched stitched regions, showing detailed overlap of AT3 and VCP ChIP-seq enrichment for the ASPS-1 cell line. (**d**) Representative tracks from highly enriched stitched regions in human ASPS tumors.

Extended Data Figure 5

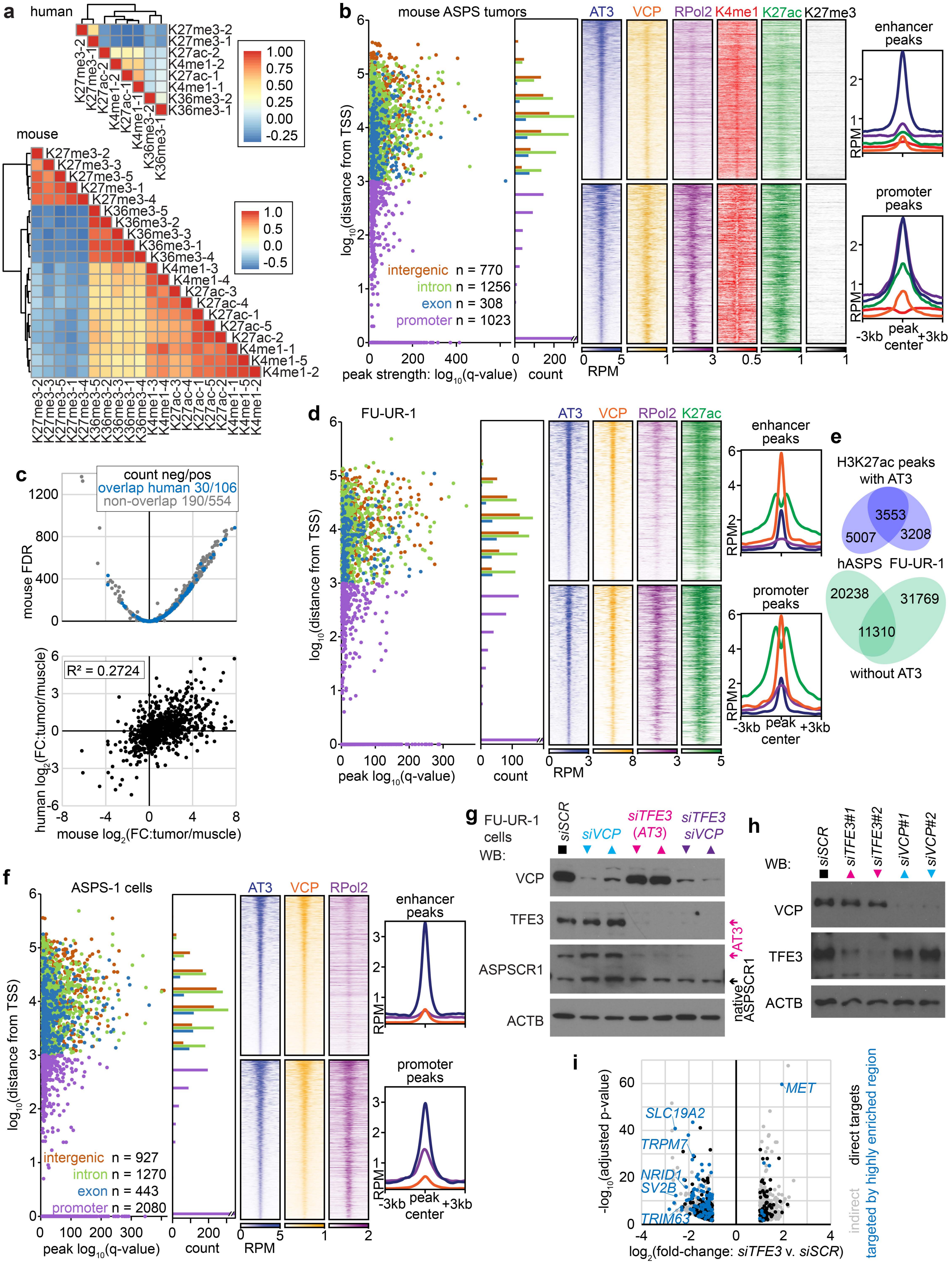

**Extended Data Figure 5. AT3:VCP interaction localizes to the promoters and enhancers of target genes.** (**a**) Pearson correlation heatmaps for the varied histone marks native ChIP-seq experiments performed on two human ASPS tumors and 5 mouse AT3-initiated ASPS tumors. (**b**) Graph and histogram of the annotations for all AT3 ChIP-seq peaks called in mouse ASPS tumors within the highly enriched regions (defined in an algorithm similar to that for human tumors in Supplemental Figure 5A). Heatmaps of reads per million (RPM) for distal (putative enhancer) and proximal promoter peaks, as well as enrichment plots for the same. (**c**) The mean differential expression of the nearest gene targets of highly enriched AT3 regions in mouse tumors compared to mouse muscle samples by RNAseq (GSE54729, FDR = false discovery rate q-value), noting the genes also annotated in human highly enriched AT3 regions and showing correlation between differential expression in human compared to mouse tumors over muscle. (**d**) Similar data for the FU-UR-1 cell line. (**e**) Venn diagrams showing overlap between human ASPS tumors and FU-UR-1 cells for H3K27ac ChIP-seq peaks genome wide that intersect with AT3 or do not. (**f**) Similar data for the ASPS-1 cell line. (**g**) Western blots (WBs) demonstrating effectiveness of knock-down by 2 different siRNAs against each of VCP, TFE3 (for AT3, as there is no native TFE3 in this male cell line), or both in human FU-UR-1 cells. (SCR = scrambled control. Triangles pointing up or down represent the siRNA#1 and siRNA#2 for each target transcript.) (**h**) WBs demonstrating modest knock-down of AT3 and VCP in ASPS-1 cells with the same siRNAs. (**i**) Differential gene expression following 48-hour siRNA depletion of AT3 in FU-UR-1 cells. Blue dots represent genes annotated by highly enriched regions (from **Extended Data Figure 4a**). Genes noted in black are those annotated by AT3 peaks generally. Gray dots denote indirect target genes, without associated AT3 ChIP-seq peaks.

Extended Data Figure 6

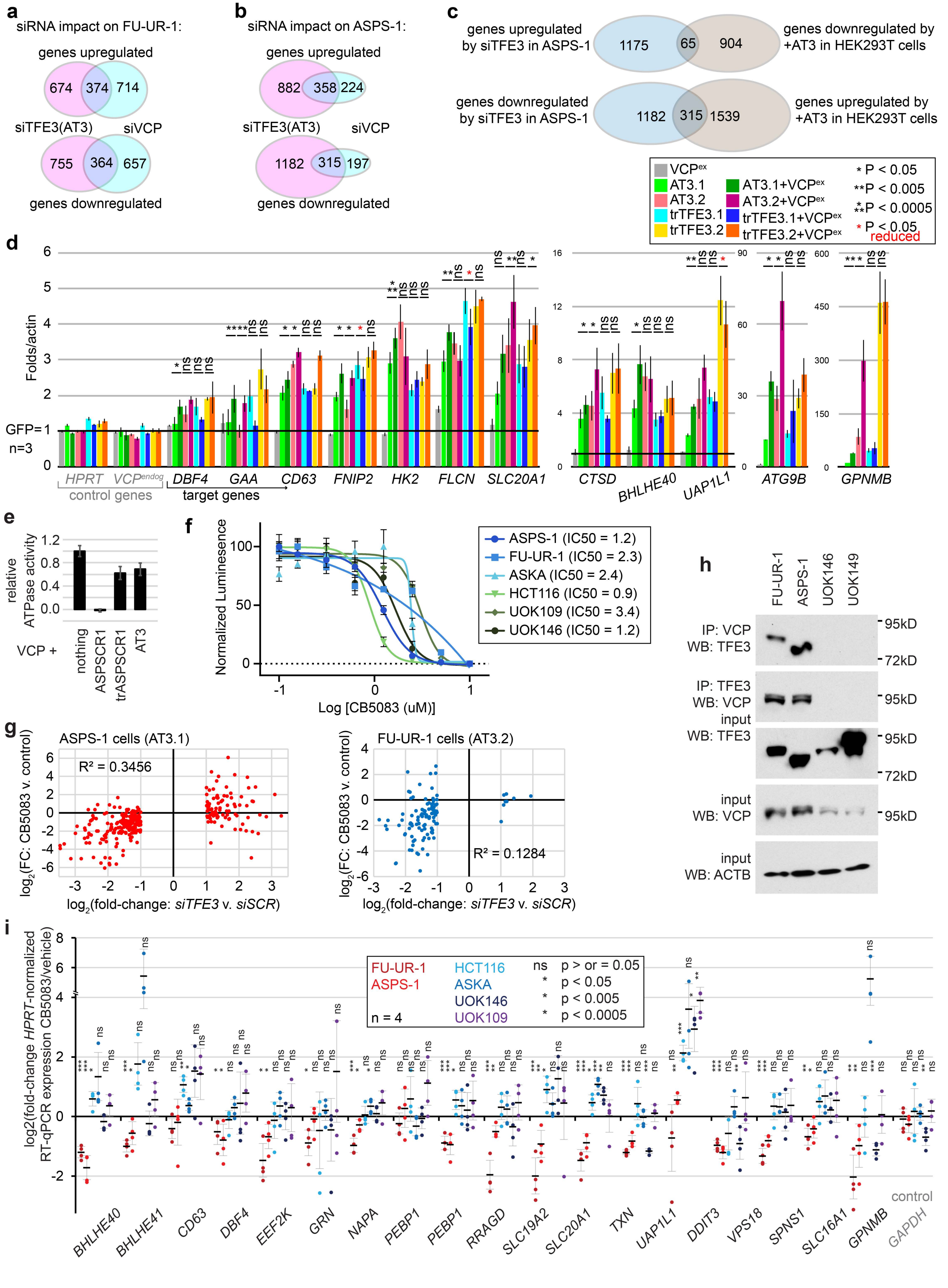

**Extended Data Figure 6. VCP presence, hexamerization, and ATPase activity impact AT3-related transcription.** (**a**) Venn diagram of overlap of whole transcriptome shifts between FU-UR-1 cells exposed for to *siTFE3* or *siVCP* (pooled from two siRNAs each and n = 3 biological replicates for each siRNA, p-adjusted < 0.05). (**b**) The same Venn diagram for ASPS-1 cells. (**c**) Overlap of significant genes changed in RNA-seq in ASPS-1 cells or HEK293T cellswith or without AT3 present. (**d**) Relative expression by RTqPCR of target genes in HEK293T cells transfected with exogenous VCP overexpression (VCPex) added to AT3.1, AT3.2, or truncated variants of TFE3 corresponding to the portions included in AT3.1 and AT3.2. Each of the statistical notations compares the addition of overexpression of VCP with the transcription factor to the transcription factor alone. (**e**) Relative ATPase activity of recombinant VCP with added recombinant ASPSCR1 (which disassembles hexamers), truncated ASPSCR1 (the portion from AT3, which does not disassemble hexamers), or AT3. (**f**) Proliferation/viability assay with Cell Titer Glo for ASPS-1, FU-UR-1, ASKA (synovial sarcoma), HCT116 (colorectal carcinoma), UOK109 (RCC with *NONO-TFE3* fusion), or UOK146 (RCC with *PRCC-TF*E3 fusion) cells showing decreasing viable cell mass after 48 hours of increasing concentrations of CB-5083, used to determine an appropriate concentration for later expression related assays, at approximately the 50 percent inhibitory concentration (n = 4). (**g**) Correlation of log-transformed fold-changes in gene expression determined by RNA-seq after CB-5083 or vehicle or pooled results of siRNA to deplete AT3 or control. These are shown for genes defined as direct targets of highly enriched regions of AT3:VCP ChIP-seq peaks and as differentially expressed at least 2-fold and significantly by depletion of AT3 over control. Each was applied for 48 hours. CB-5083 experiments had n = 2 sample size for FU-UR-1 and n = 3 for each group for ASPS-1. The siRNA samples had n = 3 for each group, but pooled results from two different siRNAs that led to depletion of AT3. (**h**) Western blots after immunoprecipitation or 10% of input for FU-UR-1 and ASPS-1 cells, as well as two RCC cell lines that express other TFE3 fusions, but do not interact with VCP. (**i**) Expression determined by RT-qPCR after exposing 6 cell lines to CB5083 for 48 hours, normalized each against HPRT expression, demonstrating the decreased expression of most AT3:VCP targets in ASPS-1 and FU-UR-1, but the opposite effect in most other cell lines, even those expressing other TFE3 fusions.

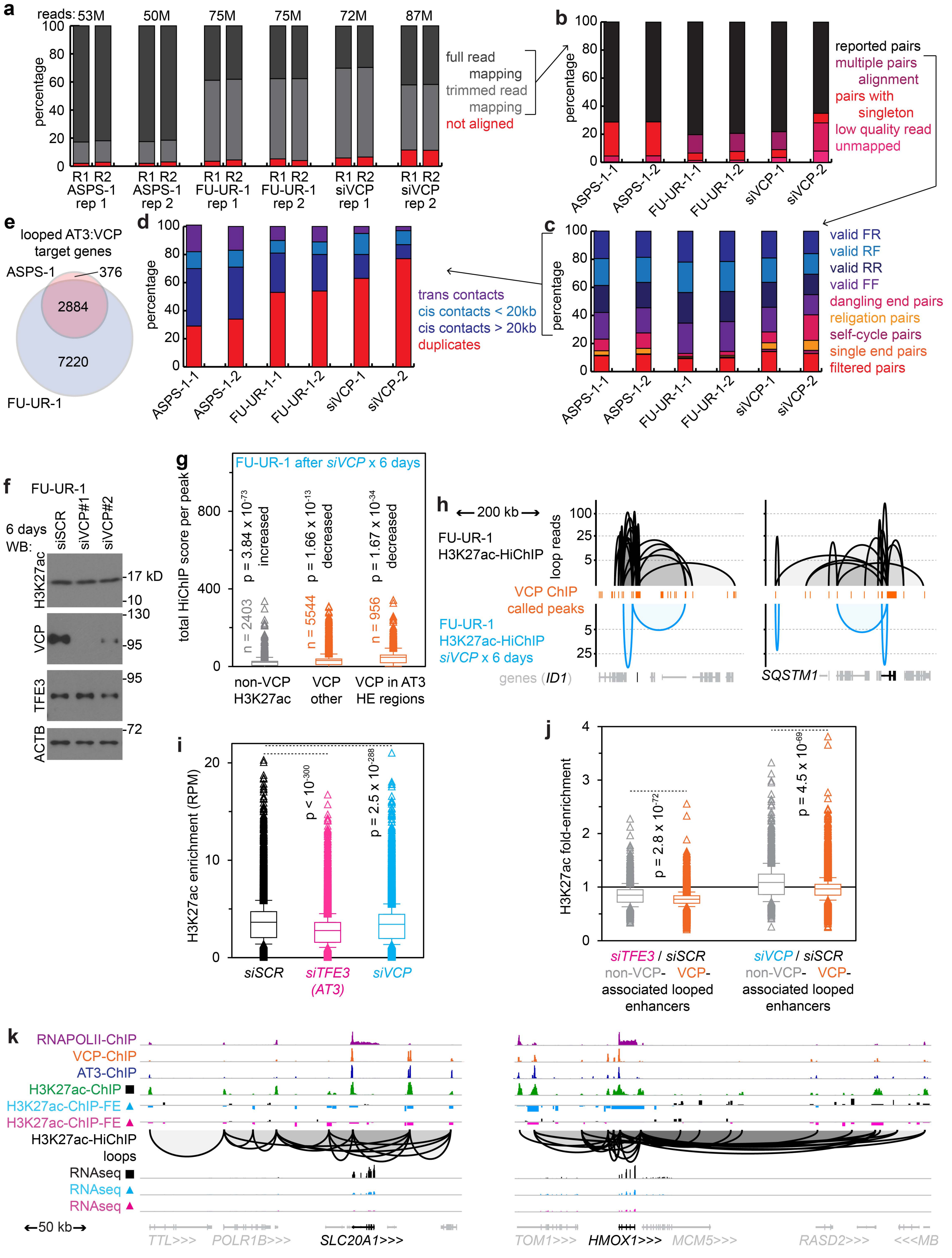
a
reads: 53M 50M 75M 75M 72M 87M
percentage
R1 R2 ASPS-1 rep 1
R1 R2 ASPS-1 rep 2
R1 R2 FU-UR-1 rep 1
R1 R2 FU-UR-1 rep 2
R1 R2 siVCP rep 1
R1 R2 siVCP rep 2
full read mapping
trimmed read mapping
not aligned
b
reported pairs
multiple pairs alignment
pairs with singleton
low quality read
unmapped
ASPS-1-1 ASPS-1-2 FU-UR-1-1 FU-UR-1-2 siVCP-1 siVCP-2
c
valid FR
valid RF
valid RR
valid FF
dangling end pairs
religation pairs
self-cycle pairs
single end pairs
filtered pairs
d
trans contacts
cis contacts < 20kb
cis contacts > 20kb
duplicates
e
looped AT3:VCP target genes
ASPS-1
376
2884
7220
FU-UR-1
f
FU-UR-1
6 days WB:
siSCR siVCP#1 siVCP#2
H3K27ac -17 kD -10
VCP -130 -95
TFE3 -95
ACTB -72
g
FU-UR-1 after siVCP x 6 days
total HiChIP score per peak
p = 3.84 x 10^-73 increased
p = 1.66 x 10^-13 decreased
p = 1.67 x 10^-34 decreased
n = 2403
n = 5544
n = 956
non-VCP H3K27ac
VCP other
VCP in AT3 HE regions
h
200 kb
FU-UR-1 H3K27ac-HiChIP
loop reads
VCP ChIP called peaks
FU-UR-1 H3K27ac-HiChIP siVCP x 6 days
genes (ID1)
SQSTM1
i
H3K27ac enrichment (RPM)
p < 10^-300
p = 2.5 x 10^-288
siSCR
siTFE3 (AT3)
siVCP
j
H3K27ac fold-enrichment
p = 2.8 x 10^-72
p = 4.5 x 10^-69
siTFE3 / siSCR
non-VCP- VCP-associated looped enhancers
siVCP / siSCR
non-VCP- VCP-associated looped enhancers
k
RNAPOLII-ChIP
VCP-ChIP
AT3-ChIP
H3K27ac-ChIP ■
H3K27ac-ChIP-FE ▲
H3K27ac-ChIP-FE ▲
H3K27ac-HiChIP loops
RNAseq ■
RNAseq ▲
RNAseq ▲
50 kb
TTL>>> POLR1B>>> SLC20A1>>>
TOM1>>> HMOX1>>> MCM5>>> RASD2>>> <<<MB

**Extended Data Figure 7. Chromatin conformation in the form of enhancer loops depend on the AT3:VCP interaction.** (**a**) Quality reports for six H3K27ac-HiChIP experiments, demonstrating the percentage and number of reads mapped per R1 and R2 tag as full reads or trimmed reads, as well as (**b**) the percentage of those read pairs that were reported or failed for the listed reasons. (**c**) The percentage of reported reads that oriented forward (F) or reverse (R), among valid pairs versus failed pairing reasons. (**d**) The distribution of trans, short or long cis, and duplicate (filtered out) contacts from the valid pairs of reported reads, per HiChIP. (**e**) Venn diagram depicting the overlap between looped target genes from **Fig. 6a-b** in the two models. (**f**) Western blots (WB) of H3K27ac and VCP, as well as controls of AT3 (TFE3) and ACTB, after prolonged exposure of FU-UR-1 cells to siSCR control or either of two siVCPs. (**g**) Box plots of the number of HiChIP loop reads per peak in FU-UR-1 cells exposed to siVCP for 6 days, by categories defined in and compared to a (2-tailed heteroschedastic t-test p-values). (**h**) H3K27ac-HiChIP loops in baseline FU-UR-1 cells or after 6 days of siVCP, with reference called peak positions for VCP ChIP. (**i**) Box plots (mean, 25th to 75th percentile, +/- standard deviation, and individual outliers, 2-tailed heteroschedastic t-test p-values) comparing H3K27ac ChIP-seq enrichment (reads per million, RPM) after 48 hours of exposure to siRNAs targeting scrambled control, TFE3 (AT3), or VCP, in FU-UR-1 cells (n = 3 for each of 2 siRNAs for each target, p-value represents a 2-tailed paired t-test showing reduced enrichment after siTFE3 or siVCP). (**j**) Box plots comparing all non-VCP-associated H3K27ac HiChIP loops to all VCP-peak-associated H3K27ac HiChIP loops for fold-enrichment for H3K27ac ChIP-seq after knock-down of AT3 or VCP, each compared to scrambled control. (**k**) Example tracks of H3K27ac ChIP-seq after knock-down of VCP or AT3 by siRNA or scrambled control. Track scales are matched between all the RNA-seq and H3K27ac-fold enrichments (FE). Autoscaling rendered ranges of RNAPOL2: 0 to 9, VCP: 0 to 15, AT3: 0 to 9, H3K27ac: 0 to 11, H3K27ac-FE: -1 to +1. HiChIP and RNA-seq read depths (both graphed on logarithmic scale) range up to 39 and 17 for HMOX1, and 32 and 16 for SLC20A1, each respectively.

Extended Data Figure 8

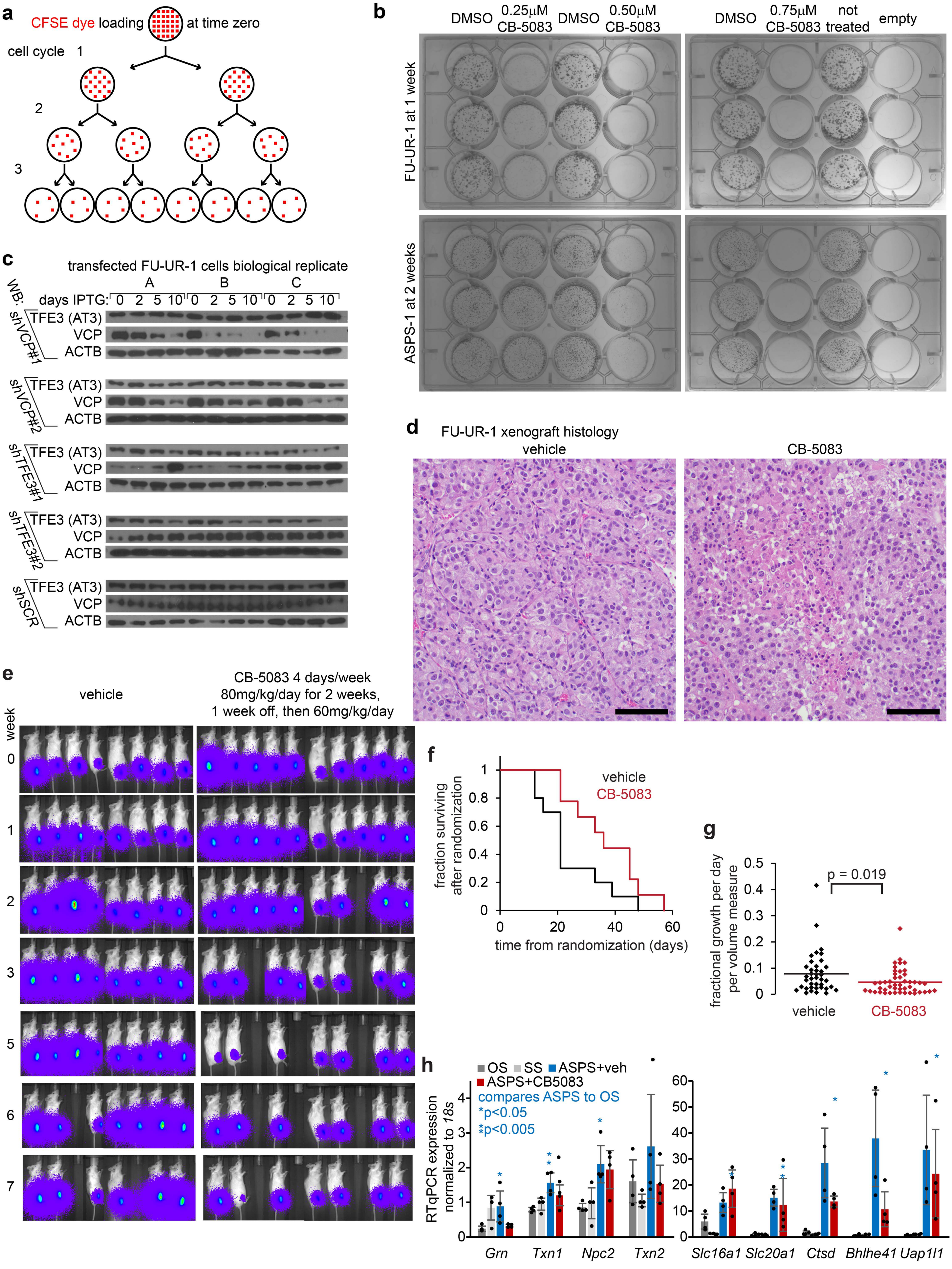

**Extended Data Figure 8. AT3:VCP is a targetable functional dependency in cancer cells**. (**a**) Schematic of CSFE dye depletion as a measure of proliferation. (**b**) Triplicate colony formation assays in FU-UR-1 and ASPS-1 cells in increasing concentrations of CB-5083 versus control DMSO to the left of each. (**c**) WBs demonstrating the modest depletion of AT3 or VCP by application of IPTG-inducible lentiviral vectors for shRNAs for the same sequences as siRNAs #1 and #2 for each target. These WBs represent whole cell lysates of FU-UR-1 cells after infection, selection, and development of 3 biological replicates for each shRNA. Each population of cells presented is after no IPTG or 2, 5, or 10 days of IPTG in culture. (**d**) Photomicrograph of hematoxylin and eosin stained sections of tumors that developed from FU-UR-1 xenografted cells into NRG mice, subsequently treated with CB-5083 or control vehicle (bar = 100mm.) (**e**) Raw luciferase FLUX data from mice implanted with an ASPS patient-derived xenograft, treated with CB-5083 or vehicle control. (**f**) Kaplan-Meier survival plots of mice randomized to treatment with vehicle (n = 10) or 50mg/kg daily 4 days per week CB-5083 by oral gavage (n = 9). (Mean time to morbidity and standard deviation 24.6 ± 11.9 days compared to 36.4 ± 12.4, respectively; two-tailed homoschedastic t-test, p = 0.034) (**g**) Fractional growth rate per day calculated from measurements at each weekly imaging for each tumor (n = 47, CB-5083; n = 39, vehicle; 2-tailed t-test p-value noted). (**h**) RT-qPCR for target genes (defined as being mouse AT3-ChIP-seq targeted and having reduced expression of homologues in both FU-UR-1 and ASPS-1 human cells upon siRNA depletion of AT3) in control tumors (OS = osteosarcoma, SS = synovial sarcoma) or genetically induced mouse ASPS tumors treated with vehicle or CB-5083 for 50mg/kg for 4 days prior to harvest. (n = 4 tumors per group per gene; homoschedastic 2-tailed t-test p-values noted, comparing vehicle-treated ASPS tumors to OS.)

**Extended Data Table 1. Enrichr Reactome pathway analysis for Fig. 4h.**

| Reactome | Overlap | p-value | Odds Ratio | Combined Score |
|---|---|---|---|---|
| ASPS-1 AT3 direct target genes | | | | |
| Activation Of Pre-Replicative Complex R-HSA-68962 | 6 out of 32 | 5.84E-07 | 24.15 | 346.65 |
| Cell Cycle, Mitotic R-HSA-69278 | 19 out of 523 | 9.42E-07 | 4.13 | 57.36 |
| Cell Cycle R-HSA-1640170 | 20 out of 654 | 6.60E-06 | 3.46 | 41.22 |
| DNA Strand Elongation R-HSA-69190 | 5 out of 31 | 1.16E-05 | 20.02 | 227.56 |
| Activation Of ATR In Response To Replication Stress R-HSA-176187 | 5 out of 36 | 2.47E-05 | 16.79 | 178.09 |
| G1/S Transition R-HSA-69206 | 8 out of 129 | 3.94E-05 | 6.96 | 70.58 |
| Cell Cycle Checkpoints R-HSA-69620 | 11 out of 271 | 7.47E-05 | 4.49 | 42.70 |
| Mitotic G1 Phase And G1/S Transition R-HSA-453279 | 8 out of 147 | 9.92E-05 | 6.05 | 55.79 |
| DNA Replication R-HSA-69306 | 8 out of 155 | 1.43E-04 | 5.72 | 50.63 |
| Synthesis Of DNA R-HSA-69239 | 7 out of 119 | 1.68E-04 | 6.55 | 56.91 |
| G1/S-Specific Transcription R-HSA-69205 | 4 out of 29 | 1.72E-04 | 16.57 | 143.63 |
| Unattached Kinetochores Signal Amplification Via A MAD2 Inhibitory Signal R-HSA-141444 | 6 out of 93 | 3.00E-04 | 7.20 | 58.36 |
| Removal Of Flap Intermediate R-HSA-69166 | 3 out of 14 | 3.07E-04 | 28.12 | 227.43 |
| EML4 And NUDC In Mitotic Spindle Formation R-HSA-9648025 | 6 out of 97 | 3.77E-04 | 6.88 | 54.22 |
| Processive Synthesis On Lagging Strand R-HSA-69183 | 3 out of 15 | 3.81E-04 | 25.77 | 202.90 |
| Mitotic Anaphase R-HSA-68882 | 9 out of 232 | 4.70E-04 | 4.25 | 32.56 |
| Mitotic Metaphase And Anaphase R-HSA-2555396 | 9 out of 233 | 4.85E-04 | 4.23 | 32.28 |
| Mitotic Prometaphase R-HSA-68877 | 8 out of 186 | 4.92E-04 | 4.72 | 35.93 |
| Resolution Of Sister Chromatid Cohesion R-HSA-2500257 | 6 out of 106 | 6.05E-04 | 6.26 | 46.36 |
| Mitotic Spindle Checkpoint R-HSA-69618 | 6 out of 110 | 7.35E-04 | 6.01 | 43.39 |
| Metabolism Of Lipids R-HSA-556833 | 17 out of 732 | 8.56E-04 | 2.55 | 18.01 |
| Lagging Strand Synthesis R-HSA-69186 | 3 out of 20 | 9.21E-04 | 18.19 | 127.14 |
| S Phase R-HSA-69242 | 7 out of 161 | 0.001036 | 4.75 | 32.65 |
| RHO GTPases Activate Formins R-HSA-5663220 | 6 out of 119 | 0.001109 | 5.53 | 37.64 |
| Separation Of Sister Chromatids R-HSA-2467813 | 7 out of 170 | 0.001420 | 4.49 | 29.42 |
| DNA Replication Pre-Initiation R-HSA-69002 | 6 out of 127 | 0.001549 | 5.16 | 33.41 |
| Fatty Acid Metabolism R-HSA-8978868 | 7 out of 173 | 0.001570 | 4.41 | 28.44 |
| CREB1 Phosphorylation Thru NMDA Receptor-Mediated Activation Of RAS Signaling R-HSA-442742 | 3 out of 27 | 0.002247 | 12.88 | 78.54 |

| | | | | |
|---|---|---|---|---|
| Metabolism R-HSA-1430728 | 33 out of 2049 | 0.002696 | 1.80 | 10.63 |
| FOXO-mediated Transcription Of Oxidative Stress, Metabolic And Neuronal Genes R-HSA-9615017 | 3 out of 29 | 0.002767 | 11.89 | 70.01 |
| Post NMDA Receptor Activation Events R-HSA-438064 | 4 out of 61 | 0.002964 | 7.26 | 42.24 |
| Nuclear Events (Kinase And Transcription Factor Activation) R-HSA-198725 | 4 out of 61 | 0.002964 | 7.26 | 42.24 |
| NR1H2 And NR1H3 Regulate Gene Expression Linked To Lipogenesis R-HSA-9029558 | 2 out of 9 | 0.003255 | 29.31 | 167.87 |
| Telomere C-strand (Lagging Strand) Synthesis R-HSA-174417 | 3 out of 33 | 0.004015 | 10.30 | 56.83 |
| Activation Of PPARGC1A (PGC-1alpha) By Phosphorylation R-HSA-2151209 | 2 out of 10 | 0.004042 | 25.64 | 141.32 |
| Creatine Metabolism R-HSA-71288 | 2 out of 10 | 0.004042 | 25.64 | 141.32 |
| M Phase R-HSA-68886 | 10 out of 380 | 0.004241 | 2.84 | 15.51 |
| Orc1 Removal From Chromatin R-HSA-68949 | 4 out of 69 | 0.004623 | 6.36 | 34.20 |
| Pyrimidine Salvage R-HSA-73614 | 2 out of 11 | 0.004909 | 22.79 | 121.18 |
| Unwinding Of DNA R-HSA-176974 | 2 out of 11 | 0.004909 | 22.79 | 121.18 |
| RHO GTPase Effectors R-HSA-195258 | 8 out of 269 | 0.005003 | 3.20 | 16.97 |
| Signaling By NTRK1 (TRKA) R-HSA-187037 | 5 out of 114 | 0.005215 | 4.76 | 24.99 |
| Activation Of NMDA Receptors And Postsynaptic Events R-HSA-442755 | 4 out of 74 | 0.005927 | 5.90 | 30.28 |
| NGF-stimulated Transcription R-HSA-9031628 | 3 out of 39 | 0.006444 | 8.58 | 43.28 |
| Kinesins R-HSA-983189 | 3 out of 42 | 0.007924 | 7.92 | 38.31 |
| Leading Strand Synthesis R-HSA-69109 | 2 out of 14 | 0.007968 | 17.09 | 82.60 |
| Signaling By NTRKs R-HSA-166520 | 5 out of 132 | 0.009557 | 4.08 | 18.96 |
| SUMOylation Of DNA Replication Proteins R-HSA-4615885 | 3 out of 45 | 0.009589 | 7.35 | 34.17 |
| Cell Surface Interactions At Vascular Wall R-HSA-202733 | 5 out of 134 | 0.010157 | 4.01 | 18.42 |
| Hemostasis R-HSA-109582 | 12 out of 576 | 0.011128 | 2.24 | 10.06 |
| Switching Of Origins To A Post-Replicative State R-HSA-69052 | 4 out of 90 | 0.011675 | 4.80 | 21.37 |
| Removal Of Flap Intermediate From C-strand R-HSA-174437 | 2 out of 17 | 0.011682 | 13.67 | 60.84 |
| Extension Of Telomeres R-HSA-180786 | 3 out of 50 | 0.012784 | 6.57 | 28.64 |
| Processive Synthesis On C-strand Of Telomere R-HSA-174414 | 2 out of 19 | 0.014502 | 12.06 | 51.06 |
| Ras Activation Upon Ca2+ Influx Thru NMDA Receptor R-HSA-442982 | 2 out of 19 | 0.014502 | 12.06 | 51.06 |
| Transcription Of E2F Targets Under Negative Control By DREAM Complex R-HSA-1362277 | 2 out of 19 | 0.014502 | 12.06 | 51.06 |
| G2/M Checkpoints R-HSA-69481 | 5 out of 148 | 0.015093 | 3.62 | 15.17 |

| | | | | |
|---|---|---|---|---|
| Regulation Of Cholesterol Biosynthesis By SREBP (SREBF) R-HSA-1655829 | 3 out of 55 | 0.016518 | 5.94 | 24.35 |
| PCNA-Dependent Long Patch Base Excision Repair R-HSA-5651801 | 2 out of 21 | 0.017584 | 10.79 | 43.61 |
| Transcriptional Regulation By MECP2 R-HSA-8986944 | 3 out of 60 | 0.020799 | 5.41 | 20.97 |
| Nucleotide Salvage R-HSA-8956321 | 2 out of 23 | 0.020917 | 9.76 | 37.75 |
| Post-chaperonin Tubulin Folding Pathway R-HSA-389977 | 2 out of 23 | 0.020917 | 9.76 | 37.75 |
| Assembly Of Pre-Replicative Complex R-HSA-68867 | 4 out of 110 | 0.022729 | 3.89 | 14.73 |
| Polymerase Switching On C-strand Of Telomere R-HSA-174411 | 2 out of 25 | 0.024490 | 8.91 | 33.06 |
| Resolution Of AP Sites Via Multiple-Nucleotide Patch Replacement Pathway R-HSA-110373 | 2 out of 25 | 0.024490 | 8.91 | 33.06 |
| FOXO-mediated Transcription R-HSA-9614085 | 3 out of 65 | 0.025633 | 4.98 | 18.23 |
| RNA Polymerase III Transcription Initiation From Type 2 Promoter R-HSA-76066 | 2 out of 26 | 0.026364 | 8.54 | 31.05 |
| Formation Of Tubulin Folding Intermediates By CCT/TriC R-HSA-389960 | 2 out of 26 | 0.026364 | 8.54 | 31.05 |
| HDR Thru Homologous Recombination (HRR) R-HSA-5685942 | 3 out of 67 | 0.027720 | 4.82 | 17.28 |
| SUMOylation R-HSA-2990846 | 5 out of 174 | 0.028053 | 3.06 | 10.93 |
| RNA Polymerase III Transcription Initiation From Type 1 Promoter R-HSA-76061 | 2 out of 27 | 0.028293 | 8.20 | 29.23 |
| Effects Of PIP2 Hydrolysis R-HSA-114508 | 2 out of 27 | 0.028293 | 8.20 | 29.23 |
| G0 And Early G1 R-HSA-1538133 | 2 out of 27 | 0.028293 | 8.20 | 29.23 |
| Chromosome Maintenance R-HSA-73886 | 4 out of 118 | 0.028473 | 3.62 | 12.87 |
| Homology Directed Repair R-HSA-5693538 | 4 out of 119 | 0.029247 | 3.59 | 12.66 |
| Prefoldin Mediated Transfer Of Substrate To CCT/TriC R-HSA-389957 | 2 out of 28 | 0.030277 | 7.88 | 27.57 |
| Energy Dependent Regulation Of mTOR By LKB1-AMPK R-HSA-380972 | 2 out of 29 | 0.032315 | 7.59 | 26.05 |
| Membrane Trafficking R-HSA-199991 | 11 out of 599 | 0.033330 | 1.95 | 6.65 |
| Regulation Of MECP2 Expression And Activity R-HSA-9022692 | 2 out of 31 | 0.036547 | 7.07 | 23.38 |
| Sealing Of Nuclear Envelope (NE) By ESCRT-III R-HSA-9668328 | 2 out of 31 | 0.036547 | 7.07 | 23.38 |
| Gluconeogenesis R-HSA-70263 | 2 out of 32 | 0.038738 | 6.83 | 22.21 |
| COPI-dependent Golgi-to-ER Retrograde Traffic R-HSA-6811434 | 3 out of 78 | 0.040764 | 4.11 | 13.15 |
| Cooperation Of Prefoldin And TriC/CCT In Actin And Tubulin Folding R-HSA-389958 | 2 out of 33 | 0.040978 | 6.61 | 21.12 |

| | | | | |
|---|---|---|---|---|
| Factors Involved In Megakaryocyte Development And Platelet Production R-HSA-983231 | 4 out of 136 | 0.044301 | 3.12 | 9.73 |
| TP53 Regulates Metabolic Genes R-HSA-5628897 | 3 out of 81 | 0.044772 | 3.95 | 12.27 |
| RNA Polymerase III Transcription Initiation R-HSA-76046 | 2 out of 35 | 0.045600 | 6.21 | 19.17 |
| Arachidonate Production From DAG R-HSA-426048 | 1 out of 5 | 0.047813 | 25.52 | 77.58 |
| Regulation Of HMOX1 Expression And Activity R-HSA-9707587 | 1 out of 5 | 0.047813 | 25.52 | 77.58 |
| Sodium-coupled Phosphate Cotransporters R-HSA-427652 | 1 out of 5 | 0.047813 | 25.52 | 77.58 |
| Sulfide Oxidation To Sulfate R-HSA-1614517 | 1 out of 5 | 0.047813 | 25.52 | 77.58 |
| HSF1-dependent Transactivation R-HSA-3371571 | 2 out of 36 | 0.047980 | 6.03 | 18.30 |
| Mitochondrial Fatty Acid Beta-Oxidation R-HSA-77289 | 2 out of 36 | 0.047980 | 6.03 | 18.30 |
| Vesicle-mediated Transport R-HSA-5653656 | 11 out of 637 | 0.048215 | 1.83 | 5.55 |
| Transcriptional Regulation Of White Adipocyte Differentiation R-HSA-381340 | 3 out of 84 | 0.048968 | 3.80 | 11.48 |
| FU-UR-1 AT3 direct target genes | | | | |
| Cell Cycle, Mitotic R-HSA-69278 | 8 out of 523 | 0.004170 | 3.346192017 | 18.33655029 |
| Circadian Clock R-HSA-400253 | 3 out of 69 | 0.004730 | 9.490909091 | 50.81218251 |
| Mitotic Anaphase R-HSA-68882 | 5 out of 232 | 0.005703 | 4.659892947 | 24.07688423 |
| Mitotic Metaphase And Anaphase R-HSA-2555396 | 5 out of 233 | 0.005805 | 4.639219015 | 23.88711048 |
| mTORC1-mediated Signaling R-HSA-166208 | 2 out of 24 | 0.006114 | 18.82575758 | 95.95691213 |
| Fertilization R-HSA-1187000 | 2 out of 25 | 0.006625 | 18.00634058 | 90.3361026 |
| Basigin Interactions R-HSA-210991 | 2 out of 25 | 0.006625 | 18.00634058 | 90.3361026 |
| BMAL1:CLOCK,NPAS2 Activates Circadian Gene Expression R-HSA-1368108 | 2 out of 27 | 0.007702 | 16.56416667 | 80.60547813 |
| RAB GEFs Exchange GTP For GDP On RABs R-HSA-8876198 | 3 out of 89 | 0.009557 | 7.276376989 | 33.83853874 |
| Separation Of Sister Chromatids R-HSA-2467813 | 4 out of 170 | 0.009802 | 5.059215586 | 23.39980205 |
| Activation Of Pre-Replicative Complex R-HSA-68962 | 2 out of 32 | 0.010713 | 13.8 | 62.60139233 |
| Unattached Kinetochores Signal Amplification Via A MAD2 Inhibitory Signal R-HSA-141444 | 3 out of 93 | 0.010769 | 6.951578947 | 31.49840393 |
| VEGFA-VEGFR2 Pathway R-HSA-4420097 | 3 out of 93 | 0.010769 | 6.951578947 | 31.49840393 |
| EML4 And NUDC In Mitotic Spindle Formation R-HSA-9648025 | 3 out of 97 | 0.012065 | 6.654423292 | 29.39544938 |
| Metabolism Of Vitamins And Cofactors R-HSA-196854 | 4 out of 186 | 0.013283 | 4.61070844 | 19.92397145 |
| Signaling By VEGF R-HSA-194138 | 3 out of 102 | 0.013807 | 6.316746411 | 27.0520303 |

| | | | | |
|---|---|---|---|---|
| Cell Cycle R-HSA-1640170 | 8 out of 654 | 0.014994 | 2.649604403 | 11.12862288 |
| Resolution Of Sister Chromatid Cohesion R-HSA-2500257 | 3 out of 106 | 0.015298 | 6.070209504 | 25.37370356 |
| Mitotic Spindle Checkpoint R-HSA-69618 | 3 out of 110 | 0.016877 | 5.842105263 | 23.84648518 |
| MTOR Signaling R-HSA-165159 | 2 out of 41 | 0.017213 | 10.61057692 | 43.10110621 |
| Reproduction R-HSA-1474165 | 3 out of 113 | 0.018118 | 5.681913876 | 22.78922929 |
| Heme Signaling R-HSA-9707616 | 2 out of 45 | 0.020521 | 9.621608527 | 37.39261943 |
| RHO GTPases Activate Formins R-HSA-5663220 | 3 out of 119 | 0.020751 | 5.386388385 | 20.87320759 |
| Rab Regulation Of Trafficking R-HSA-9007101 | 3 out of 122 | 0.022142 | 5.249800973 | 20.00330253 |
| Metabolism Of Water-Soluble Vitamins And Cofactors R-HSA-196849 | 3 out of 122 | 0.022142 | 5.249800973 | 20.00330253 |
| Hemostasis R-HSA-109582 | 7 out of 576 | 0.023111 | 2.613627146 | 9.846718886 |
| Inositol Phosphate Metabolism R-HSA-1483249 | 2 out of 48 | 0.023161 | 8.992753623 | 33.86022788 |
| Arachidonate Production From DAG R-HSA-426048 | 1 out of 5 | 0.024263 | 51.28350515 | 190.7125852 |
| Toxicity Of Botulinum Toxin Type D (botD) R-HSA-5250955 | 1 out of 5 | 0.024263 | 51.28350515 | 190.7125852 |
| Toxicity Of Botulinum Toxin Type F (botF) R-HSA-5250981 | 1 out of 5 | 0.024263 | 51.28350515 | 190.7125852 |
| Vitamin B1 (Thiamin) Metabolism R-HSA-196819 | 1 out of 5 | 0.024263 | 51.28350515 | 190.7125852 |
| Regulation Of HMOX1 Expression And Activity R-HSA-9707587 | 1 out of 5 | 0.024263 | 51.28350515 | 190.7125852 |
| Drug-mediated Inhibition Of CDK4/CDK6 Activity R-HSA-9754119 | 1 out of 5 | 0.024263 | 51.28350515 | 190.7125852 |
| G1/S Transition R-HSA-69206 | 3 out of 129 | 0.025581 | 4.956390977 | 18.16966644 |
| Cell Surface Interactions At Vascular Wall R-HSA-202733 | 3 out of 134 | 0.028204 | 4.766010446 | 17.00657294 |
| PTK6 Regulates Cell Cycle R-HSA-8849470 | 1 out of 6 | 0.029046 | 41.02474227 | 145.1819826 |
| CD22 Mediated BCR Regulation R-HSA-5690714 | 1 out of 6 | 0.029046 | 41.02474227 | 145.1819826 |
| Vitamin B2 (Riboflavin) Metabolism R-HSA-196843 | 1 out of 7 | 0.033805 | 34.18556701 | 115.7918426 |
| Zinc Efflux And Compartmentalization By SLC30 Family R-HSA-435368 | 1 out of 7 | 0.033805 | 34.18556701 | 115.7918426 |
| Constitutive Signaling By NOTCH1 t(7;9)(NOTCH1:M1580 K2555) Translocation Mutant R-HSA-2660826 | 1 out of 7 | 0.033805 | 34.18556701 | 115.7918426 |
| Mitotic G1 Phase And G1/S Transition R-HSA-453279 | 3 out of 147 | 0.035666 | 4.332894737 | 14.44395145 |
| RAB Geranylgeranylation R-HSA-8873719 | 2 out of 62 | 0.037157 | 6.889583333 | 22.68475283 |
| Axonal Growth Inhibition (RHOA Activation) R-HSA-193634 | 1 out of 8 | 0.038541 | 29.30044183 | 95.40343353 |

| | | | | |
|---|---|---|---|---|
| VEGF Binds To VEGFR Leading To Receptor Dimerization R-HSA-195399 | 1 out of 8 | 0.038541 | 29.30044183 | 95.40343353 |
| M Phase R-HSA-68886 | 5 out of 380 | 0.039066 | 2.799569892 | 9.077637199 |
| Sperm Motility And Taxes R-HSA-1300642 | 1 out of 9 | 0.043254 | 25.63659794 | 80.51615206 |
| Activation Of PUMA And Translocation To Mitochondria R-HSA-139915 | 1 out of 9 | 0.043254 | 25.63659794 | 80.51615206 |
| p75NTR Regulates Axonogenesis R-HSA-193697 | 1 out of 9 | 0.043254 | 25.63659794 | 80.51615206 |
| RHO GTPase Effectors R-HSA-195258 | 4 out of 269 | 0.043341 | 3.153271778 | 9.897072334 |
| Cell Cycle Checkpoints R-HSA-69620 | 4 out of 271 | 0.044328 | 3.129333015 | 9.751413208 |
| S Phase R-HSA-69242 | 3 out of 161 | 0.044726 | 3.946169221 | 12.26149807 |
| Orc1 Removal From Chromatin R-HSA-68949 | 2 out of 69 | 0.045098 | 6.167599502 | 19.11288794 |
| Synthesis Of Pyrophosphates In Cytosol R-HSA-1855167 | 1 out of 10 | 0.047944 | 22.78694158 | 69.22044886 |
| Glycoprotein Hormones R-HSA-209822 | 1 out of 10 | 0.047944 | 22.78694158 | 69.22044886 |
| HuR (ELAVL1) Binds And Stabilizes mRNA R-HSA-450520 | 1 out of 10 | 0.047944 | 22.78694158 | 69.22044886 |
| Interaction With Cumulus Cells And Zona Pellucida R-HSA-2534343 | 1 out of 10 | 0.047944 | 22.78694158 | 69.22044886 |
| Neurotoxicity Of Clostridium Toxins R-HSA-168799 | 1 out of 10 | 0.047944 | 22.78694158 | 69.22044886 |

**Extended Data Table 2. Contribution coefficients for genes in PCA plot in Fig. 6g**

| Gene symbol | PC1 coefficient | PC2 coefficient | PC1 coefficient absolute value |
|---|---|---|---|
| *INHBE* | -0.370177426 | -0.301028375 | 0.370177426 |
| *NEU1* | -0.23756408 | 0.114892569 | 0.23756408 |
| *MIOX* | -0.226201812 | 0.182219044 | 0.226201812 |
| *CDK4* | -0.221605632 | 0.078480517 | 0.221605632 |
| *MET* | 0.204242236 | -0.190206091 | 0.204242236 |
| *BHLHE41* | -0.202009127 | 0.090222659 | 0.202009127 |
| *ADM2* | -0.187311597 | -0.128412565 | 0.187311597 |
| *RAB3IL1* | -0.184383075 | -0.050373303 | 0.184383075 |
| *CPVL* | -0.181193723 | 0.076631733 | 0.181193723 |
| *CREB3L1* | -0.171689357 | -0.040614838 | 0.171689357 |
| *RAB32* | -0.169800702 | 0.06692255 | 0.169800702 |
| *RALGDS* | -0.163491216 | -0.006658371 | 0.163491216 |
| *CPEB1* | -0.154801813 | -0.050176279 | 0.154801813 |
| *CUTA* | -0.147634547 | 0.125545792 | 0.147634547 |
| *ID1* | -0.144214803 | 0.144768984 | 0.144214803 |
| *MYEOV* | -0.141433162 | 0.0334767 | 0.141433162 |
| *BAIAP2* | -0.137976166 | -0.058833466 | 0.137976166 |
| *DNAAF5* | -0.136756659 | -0.014085813 | 0.136756659 |
| *WWC1* | 0.136120743 | 0.044662359 | 0.136120743 |
| *GDF15* | -0.135570978 | -0.372389726 | 0.135570978 |
| *BIRC7* | -0.134164185 | 0.072546278 | 0.134164185 |
| *PATL1* | -0.132084036 | 0.024830057 | 0.132084036 |
| *BHLHE40* | -0.126110289 | 0.054890496 | 0.126110289 |
| *PRKAG2* | -0.118239919 | -0.082576269 | 0.118239919 |
| *CHN2* | -0.114306268 | -0.021319696 | 0.114306268 |
| *CAMKK1* | -0.108162896 | 0.040990243 | 0.108162896 |
| *ATP6V1B2* | -0.105905979 | -0.007316181 | 0.105905979 |
| *ITPR1* | -0.099972804 | -0.001607185 | 0.099972804 |
| *SLC25A13* | -0.099517538 | 0.09431633 | 0.099517538 |
| *PLCD3* | 0.096871775 | -0.026859817 | 0.096871775 |
| *SORBS3* | -0.095040381 | 0.080913904 | 0.095040381 |
| *ALYREF* | -0.094476586 | -0.011427483 | 0.094476586 |
| *VAC14* | -0.093091833 | 0.024137256 | 0.093091833 |
| *METTL8* | -0.087959071 | 0.130630362 | 0.087959071 |
| *PRODH* | -0.083462264 | 0.064666262 | 0.083462264 |
| *BCAR1* | 0.083397057 | -0.009385072 | 0.083397057 |
| *KCP* | 0.081180388 | 0.035898894 | 0.081180388 |
| *RHEB* | -0.075609707 | -0.085094179 | 0.075609707 |
| *ZBED6CL* | 0.074959588 | -0.016805224 | 0.074959588 |
| *KIFC3* | 0.068814944 | 0.013333436 | 0.068814944 |
| *EIF4B* | 0.068610431 | -0.049181414 | 0.068610431 |
| *ACTR3C* | -0.064483237 | -0.067032979 | 0.064483237 |
| *HNRNPF* | -0.062059255 | 0.032376444 | 0.062059255 |

| | | | |
|---|---|---|---|
| *MAFG* | -0.061336395 | -0.146295783 | 0.061336395 |
| *TRIP6* | 0.060917863 | -0.116458171 | 0.060917863 |
| *S100A2* | -0.058975353 | 0.005179431 | 0.058975353 |
| *WBP2* | -0.056809905 | -0.013018373 | 0.056809905 |
| *GAPDH* | -0.054934131 | 0.008708966 | 0.054934131 |
| *ABCC3* | 0.054168509 | 0.1205592 | 0.054168509 |
| *ACBD4* | 0.052472913 | 0.035934527 | 0.052472913 |
| *KRT80* | 0.051047281 | -0.123381685 | 0.051047281 |
| *NKAIN4* | 0.05103473 | 0.149636179 | 0.05103473 |
| *HES1* | 0.050819845 | -0.103041307 | 0.050819845 |
